# Disruption of the Homer1 coiled-coiled domain by a novel de novo human *HOMER1* variant impairs protein scaffolding, calcium signalling, and synaptogenesis

**DOI:** 10.64898/2026.08.05.741753

**Authors:** Daniel Bligh, Lisa Foa, Robert Gasperini

## Abstract

Rare de novo variants in synaptic scaffolding proteins are increasingly recognized for their roles in driving abnormal neuronal connectivity underlying conditions such as epilepsy and autism spectrum disorder (ASD). Homer1b/c, a synaptic scaffolding protein, regulates a wide suite of synaptic functions including Ca^2+^ signalling, dendritic spine morphogenesis and multiple forms of synaptic plasticity. Here we report a novel substitution mutation in the human *HOMER1* gene, *HOMER1^R297W^*, and demonstrate that Homer1b/c^R297W^ expression dominant-negative like effect on Homer1-dependent functions. In dorsal root ganglion (DRG) sensory neurons, Homer1b/c1^R297W^ impairs axonal growth cone turning to gradients of brain-derived neurotrophic factor (BDNF), a process that requires functional store-operated Ca^2+^ entry (SOCE). Accordingly, we found that SOCE was significantly blunted in both Homer1b/c^R297W^ DRG growth cones and hippocampal neuron soma. In hippocampal neurons, Homer1b/c^R297W^ lowered dendritic spine density and reduced endoplasmic reticulum infiltration into spines. Homer1b/c^R297W^ hippocampal neurons also exhibited decreased synaptic metabotropic glutamate receptor 5 (mGluR5) expression and blunted dendritic Ca²⁺ increases following group-I mGluR activation. Super resolution imaging using direct stochastic optical reconstruction microscopy (dSTORM) further demonstrated that Homer1b/c^R297W^ diminishes receptor clustering, uncoupling it from crucial binding partners including IP3R, mGluR5 and STIM1/2. Taken together, these findings highlight the importance of Homer1b/c’s tetrameric scaffolding in shaping axon guidance, dendritic spine dynamics and synaptic Ca²⁺ signalling. Disruption of these processes by Homer1b/c^R297W^ offers valuable mechanistic insights into how rare de novo variants and altered protein scaffolding can contribute to the connectivity deficits implicated in neurodevelopmental and neurological disorders.

## Introduction

Precise synaptic connectivity is essential for central nervous system (CNS) function and relies upon tightly regulated molecular signalling processes. When such signalling is disrupted, synaptic miswiring can arise, which is implicated in the pathogenesis of multiple neurological conditions including autism spectrum disorder (ASD) (Supekar et al., 2013; Lord et al., 2018; Pagani et al., 2026). A growing body of research emphasises the importance of synaptic scaffolding proteins in shaping the processes that regulate proper circuit formation during neurodevelopmental processes, including axon guidance and dendritic spinogenesis (Luo et al., 2012; Sala et al., 2015a; Soler et al., 2018; Vyas et al., 2021; Barone et al., 2026).

Previous research has identified the *HOMER1* gene, which encodes the synaptic scaffolding protein Homer1, as an autism-risk gene (Kelleher III et al., 2012). *HOMER1* produces multiple isoforms through alternative splicing, including the constitutively expressed long forms Homer1b and Homer1c, and the activity-inducible short form Homer1a (Shiraishi-Yamaguchi and Furuichi, 2007). Each isoform contains an N-terminal Ena/Vasp Homology 1 (EVH1) domain, through which Homer1 binds to ligands including inositol 1,4,5-trisphosphate receptors (IP3R) and metabotropic glutamate receptors (mGluR) (Tu et al., 1998, 1999), along with other scaffolding proteins comprising the postsynaptic density including Shank (Hayashi et al., 2009). Through its interactions with and scaffolding of such proteins via EVH1-mediated binding, Homer1 has been implicated in regulation of multiple forms of both Hebbian and non-Hebbian plasticity in vitro and in vivo (Szumlinski et al., 2006; Tappe and Kuner, 2006; Roloff et al., 2010; Gerstein et al., 2012; Gimse et al., 2018; Bockaert et al., 2021; Heavner et al., 2021; Guerrero and Turrigiano, 2025a)

Homer1b and Homer1c, collectively referred to as Homer1b/c due to their structural and functional similarity, also possess a C-terminal region comprised of a coiled-coil domain interspaced by two leucine-zipper motifs. The C-terminal region facilitates the formation of homophilic structures composed of multiple Homer proteins aligned in parallel to form homotetramers, with EVH1 domains positioned as both ends for ligand binding (Hayashi et al., 2006, 2009). These homotetramers further associate with the complementary postsynaptic scaffolding protein, Shank, to assemble higher-order scaffolding complexes (Hayashi et al., 2009). These complexes act as signalling hubs, linking group-I mGluR to endoplasmic reticulum (ER) Ca^2+^ release and downstream synaptic plasticity changes (Tu et al., 1998). Homer1b/c has also been implicated in the regulation of store-operated Ca^2+^ entry (SOCE) - a process necessary for Ca^2+^-dependent axon guidance (Mitchell et al., 2012; Pavez et al., 2019) - through interactions with both the ER-Ca^2+^ sensor STIM1 (Dionisio et al., 2015; Rao et al., 2016), transient receptor potential cation channels (TRPCs) (Yuan et al., 2003, 2012), and store-operated channel Orai1 (Jia et al., 2017). Thus, Homer1b/c coiled-coil integrity is crucial for the function of the protein, and mutations within this domain may significantly perturb synaptic function.

Reduced Homer1b/c expression is implicated in the pathogenesis of Fragile X syndrome (Giuffrida et al., 2005; Aloisi et al., 2017), and GWAS studies have separately associated *HOMER1* with ASD (Kelleher III et al., 2012). More broadly, mutations affecting other postsynaptic scaffolding proteins, particularly Shank, have been well established as contributors to ASD pathogenesis (Peça et al., 2011; Sala et al., 2015b; Zhou et al., 2016, 2019), and have been linked to aberrant Ca^2+^-dependent signalling processes through pathways also regulated by Homer1b/c (Qin et al., 2022). Yet despite the convergence of genetic and functional evidence pointing to Homer1b/c as a plausible candidate of neurodevelopmental disease risk, no study has directly tested the functional consequences of a rare, de novo Homer1b/c variant on core aspects of neuronal function including Ca^2+^ signalling.

Here we present a novel de novo human variant, *HOMER1^R297W^*, which substitutes arginine 297 for tryptophan within the coiled-coil domain of Homer1b/c. Previous structural studies show that this residue mediates salt bridges between aligned Homer1 molecules in the C1 region, stabilising dimer formation (Hayashi et al., 2009). Because tetramers form via tail-to-tail alignment of two such dimers, disruption of these C1-mediated interactions is likely to impair tetramerisation. We hypothesised that loss of positive charge at position 297 thereby would perturb dimerization, downstream tetramer assembly and thereby impair Homer1b/c function. We demonstrate here that Homer1b/c^R297W^ expression exerts a dominant-negative-like effect on Homer1-dependent processes including Ca^2+^ signalling and the scaffolding of postsynaptic protein complexes. These molecular defects correlate with both decreased dendritic-spine density and perturbed axon guidance, processes that contribute to the overall pattern of neuronal connectivity which is often aberrant in neuronal disorders. Our findings offer important insights into how scaffolding protein function can shape various aspects of neuronal function and illustrate how rare de novo variants can illuminate common molecular mechanisms underpinning disorders of synaptic connectivity.

## Materials and Methods

### Animals

All animal experiments were conducted with the approval of the University of Tasmania Animal Ethics Committee under the ethics numbers A17373 and A0025066, and in accordance with the Australian NHMRC Code of Practice for the Care and Use of Animals for Scientific Purposes. Sprague Dawley and Long Evans rats were used.

### Primary hippocampal neuron culture

Primary hippocampal neurons were prepared from E16-18 Sprague Dawley or Long Evans rat embryos. Hippocampi were dissected in ice-cold Ca^2+^/Mg^2+^-free EBSS, digested with 0.25% trypsin and 0.125% DNAse for 15 min at 37°C, and mechanically dissociated by gentle trituration. Neurons were plated on poly-L-lysine (1mg/ml) and laminin (50 µg/ml)-coated glass coverslips in Neurobasal medium supplemented with B27 (1X) and penicillin-streptomycin (100 U/ml). Cultures were maintained at 37°C in 5% CO_2_.

### Primary DRG sensory neuron culture

Primary cultures from DRG sensory neurons were prepared as previously described (Gasperini et al., 2009; Pavez et al., 2019). Briefly, thoracic DRG from E16-18 Sprague Dawley or Long Evans rat embryos were mechanically dissociated and plated onto poly-L-ornithine (1mg/ml) and laminin (50 µg/ml)-coated glass coverslips in sensory neuron media (SNM): DMEM F-12 medium 1:1, penicillin-streptomycin (100 µg/ml), N2 neural medium supplement (1%), FCS (5%) and nerve growth factor (50 ng/ml). Cultures were maintained at 37°C in 5% CO_2_ humidified incubator for 4-6hr before imaging.

### Transfection

Hippocampal and DRG sensory neurons were transfected with a rat neuron Nucleofector kit (Lonza VPG1003) and Nucleofector 2b device (Lonza AAB-1001) with 2-4µg of DNA according to the manufacturer’s instructions. Plasmids included the following: BiP-mCherry-KDEL (based on (Zurek et al., 2011). Homer1b^WT-GFP^, Homer1b^R297W-GFP^, Homer1b^WT:IRES-GFP^ and Homer1b^R297W:IRES-GFP^.

### Growth cone turning assay

Growth cone turning assays were conducted as previously described (Lohof et al., 1992; Li et al., 2005a; Wang and Poo, 2005; Gasperini et al., 2009). Briefly, micropipettes with a 1µm opening were loaded with 2.5µl of 10µg/ml BDNF. Gradients were established using a pulsatile ejection system (Picospritzer 3, Parker Hannifin) with a positive ejection pressure of 5 psi and pulses of 1 Hz. Images were acquired every 7 secs for 30 mins using NIS elements 4.5 (Nikon). Axon extension and growth cone turning angles were analysed in exported .tif files using ImageJ. Axons that failed to extend 10 µm or more over this time were excluded from analysis. Angular measurements taken from growth cones turning towards the micropipette were expressed as positive values indicative of an attractive turn and turns in the opposite direction as negative values.

### Calcium imaging

All Ca^2+^ imaging was performed using the ratiometric indicator Fura-2 AM. Neurons were loaded with 0.5 µM Fura-2AM in calcium-replete imaging buffer for 7 min at 37°C, 5% CO_2_, then washed and transferred to a temperature-controlled imaging stage (TC-324C, Warner Instruments). All imaging was performed on a Nikon Eclipse Ti microscopy with a 40x S Fluor DIC objective (NA 1.3) and Photometrics Evolve camera, with image acquisition controlled by NIS elements AR 4.5.

For SOCE recordings, neurons were imaged for 33 min with frames acquired at 5 s intervals. Following a 5.5 min baseline period in calcium-replete buffer, media was exchanged for calcium-free buffer, then calcium-free buffer containing 10 µM cyclopiazonic acid (CPA) at 10.5 min to deplete ER Ca²⁺ stores and promote CRAC channel formation. Calcium-replete buffer was restored at 20.5 min to initiate Ca²⁺ influx. SOCE amplitude was calculated as the difference between the final F_340_/F_380_ ratio recorded in calcium-free/CPA conditions and the peak ratio following reintroduction of extracellular Ca²⁺. DRG neurons were imaged >6 hrs post-plating to allow sufficient construct expression. Hippocampal neurons were imaged at 10–16 DIV, with equal numbers per condition at each timepoint.

For mGluR-dependent Ca²⁺ recordings, hippocampal neurons were imaged for 10 min at 5 s intervals. Following a 4.5 min baseline, bath application of 200 µM 3,5-DHPG in calcium-replete buffer was performed and imaging continued for a further 5.5 min. Ca²⁺ responses were measured from ROIs 40µm in length drawn around dendrites of transfected neurons.

All F_340_/F_380_ ratio data were background-corrected using a 10×10 µm cell-free ROI and exported from NIS Elements for statistical analysis. Ca²⁺ transients were defined as increases of ≥20% above baseline average persisting for no more than two frames (10 s).

### Immunocytochemistry

Cells were fixed in 4% paraformaldehyde in PBS for 10 min at room temperature, then blocked and permeabilised in 5% FCS / 0.1% Triton-X-100 in PBS for 30 min. Primary antibodies were incubated overnight at 4°C immunostaining for Homer1b/c (1:500 conventional, 1:200 dSTORM, anti-Homer1b/c, Synaptic Systems Cat# 160023, RRID:AB_2619858), GFP (1:500 conventional, 1:300 dSTORM, anti-GFP, Abcam Cat# Ab17390, RRID:AB_300798), IP_3_R (1:200 dSTORM, anti-IP_3_R, Abcam, Cat# Ab5804, RRID:AB_305124), mGluR5 (1:250 conventional, 1:200 dSTORM, anti-mGluR, Abcam Cat# ab76316, RRID:AB_1523944), STIM1 (1:200 dSTORM, anti-STIM1, Sigma-Aldrich Cat#S6072, RRID:AB_1079008), STIM2 (1:200 dSTORM, anti-STIM2, Sigma-Aldrich Cat# PRS4123, RRID:AB_1857580), mCherry (1:500 conventional, anti-mCherry, Thermo Fisher Scientific Cat# MA5-32977, RRID:AB_2802611). Secondary antibodies including anti-chicken 488 were incubated for 90 min at room temperature including alpaca 568/647 nanobodies (1:500-1000 conventional, Thermo Fisher Scientific Cat# SA5-10331, SA5-10325, SA5-10327, RRID:AB_2868378, RRID:AB_2868372, RRID:AB_2868374), Alexa Fluor 488 (1:1000 conventional/dSTORM, Thermo Fisher Scientific Cat# A-11039, RRID:AB_142924), Alexa Fluor 647 (1:300 dSTORM, Abcam Cat# Ab150171, RRID:AB_2921318), CF 680 (1:300 dSTORM, Biotium Cat# 20418). 3×10 min PBS washes were performed between each step. Coverslips were mounted in DPX (Sigma-Aldrich). Super-resolution samples were imaged in Attofluor cell chambers (Invitrogen).

### Widefield and confocal microscopy

Epifluorescence imaging was performed on a Nikon Eclipse Ti2 equipped with an iXon Life EMCCD camera (Andor) and 100× Apo TIRF objective (NA 1.49), with image acquisition in NIS Elements 4.5. Images were acquired as Z-stacks with 0.1 µm steps. Exposure settings were held constant across conditions for each channel. Confocal imaging of hippocampal neuron spines was performed on a spinning disk confocal (UltraView, PerkinElmer) with a 40× Plan Apo objective using Volocity software. Dendritic spines were quantified from 50 µm stretches of secondary or higher-order dendrites using Neurolucida 360 2021 (MBF Bioscience). mGluR5 and KDEL-mCherry fluorescence intensity within spine heads was quantified in Fiji from maximum intensity projections. Homer1b/c-mGluR5 coupling in growth cones was quantified by statistical object distance analysis (SODA) (Lagache et al., 2018) implemented in Icy bioimage software, using the "Easy SODA 2 colours; 1 image" protocol with spot detector sensitivity of 70 for both channels.

### dSTORM super-resolution microscopy

dSTORM imaging was performed on a Nikon Eclipse Ti with a 100× Apo TIRF objective (NA 1.49) and Abbelight SAFe 360 system, using NEO_liveimaging software (Abbelight). Samples were imaged in a reducing/oxidising buffer comprising 100 mM MEA-HCl, 4 µg/ml catalase and 100 µg/ml glucose oxidase, pH 7.8-8.0, refreshed every 1.5 hr. Two-dimensional spectral demixing acquisitions consisted of 15,000 frames at 0.05 ms exposure using a 640 nm laser, with illumination angle optimised per cell to achieve uniform HiLo-proximal excitation.

All localisation data were processed in Abbelight NEO_analysis software. The first 1,000 frames were excluded to ensure only stochastic photoswitching events were analysed. Spectral demixing separated anti-GFP 647 and target epitope 680 signals based on photon ratio (I1/(I1+I2)); acquisitions with indistinct ratio peaks were excluded. Cluster analysis of STIM2, mGluR5 and IP3R localisations in dendritic spine heads was performed using Voronoi tessellation within NEO_analysis, based on the SR-Tesseler method (Levet et al., 2015). For each neuron, five adjacent spine heads expressing the greatest quantity of the epitope of interest were sampled. The spatial coupling of Homer1b/c in growth cones to mGluR5 and STIM1 was analysed using the 2D-STORM SODA plugin in Icy (Lagache et al., 2018). Co-ordinate based colocalisation (CBC) analysis was performed using the CBC package within NEO_analysis. To control for substantially different labelling densities between cells, we expressed the number localisations at a given coefficient value as a percentage of the sum of all localisations across the full range of -1 to +1 CBC coefficients. In line with previous work using CBC to compare colocalisation of super-resolution resolved puncta (Harwardt et al., 2020), CBC values were filtered for positive CBC values (>0.15) representing positively colocalised interactions. The weighted mean of these values was computed per cell and compared across conditions.

### Experimental Design and Statistical Analysis

All quantitative data was collated and organised in Microsoft Excel, and all tests for significance were performed using GraphPad Prism. Statical tests comparing means between 2 groups utilised Welch’s t-test for normally distributed datasets. The Mann-Whitney U-test was used for comparisons between 2 groups with non-parametric distributions. When comparing 3 or more groups, we used one-way ANOVA with Tukey’s test for multiple comparisons to test within all groups, or one-way ANOVA with Dunnet’s test for multiple comparisons when comparing all groups against a specific control. Data normality was tested with the Shapiro-Wilk test. P-values are defined as *p = < 0.05, ** p = < 0.005, *** p = < 0.0005, **** p = < 0.00005.

## Results

### Homer1b/c^R297W^ expression disrupts Ca^2+^-dependent axon guidance in sensory neuron growth cones

To characterise the functional consequences of *HOMER1^R297,^* we first examined its effect upon axon guidance in sensory neuron growth cones, utilising an assay offering both a neurodevelopmentally relevant process and a well described Ca^2+^-dependent phenotypic readout. Although Homer1b/c has been most extensively studied as a postsynaptic scaffolding protein, it is also widely expressed in the axon and axonal growth cone. Axons in the developing nervous system are directed to target cells and achieve functional circuitry through the reception of guidance cues in the external environment, received chiefly at the filopodia tipping the distal end of the growth cone, and the motile responses of growth cones to many of these cues are Ca^2+^-dependent. Homer1b/c regulates both ER-Ca^2+^ release through IP3R and store-operated calcium entry through TRPCs (Yuan et al., 2003, 2012), key processes in several forms of Ca^2+^-dependent guidance (Li et al., 2005b; Shim et al., 2013; Pavez et al., 2019). Indeed, Homer1b/c expression has been shown to be necessary for axon guidance both in vitro (Gasperini et al., 2009) and in vivo (Foa et al., 2001). We hypothesised that Homer1b/c^R297W^ expression would impair Ca^2+^ signalling during Ca^2+^-dependent axon guidance, resulting in aberrant growth cone motile responses to a guidance cue.

To test this, the expression of Homer1 proteins in DRG sensory neurons was manipulated either through knockdown of endogenous Homer1b/c using siRNA, or overexpression of Homer1b/c^WT:IRES-GFP^ or Homer1b/c^R297W:IRES-GFP^ through expression vectors. These sensory neurons were then subjected to a well-established in vitro growth cone motility assay to characterise their motile responses to the Ca^2+^-dependent guidance cue, BDNF (Fig. 1A) (Fan and Raper, 1995; Li et al., 2005b).

**Figure 1:**
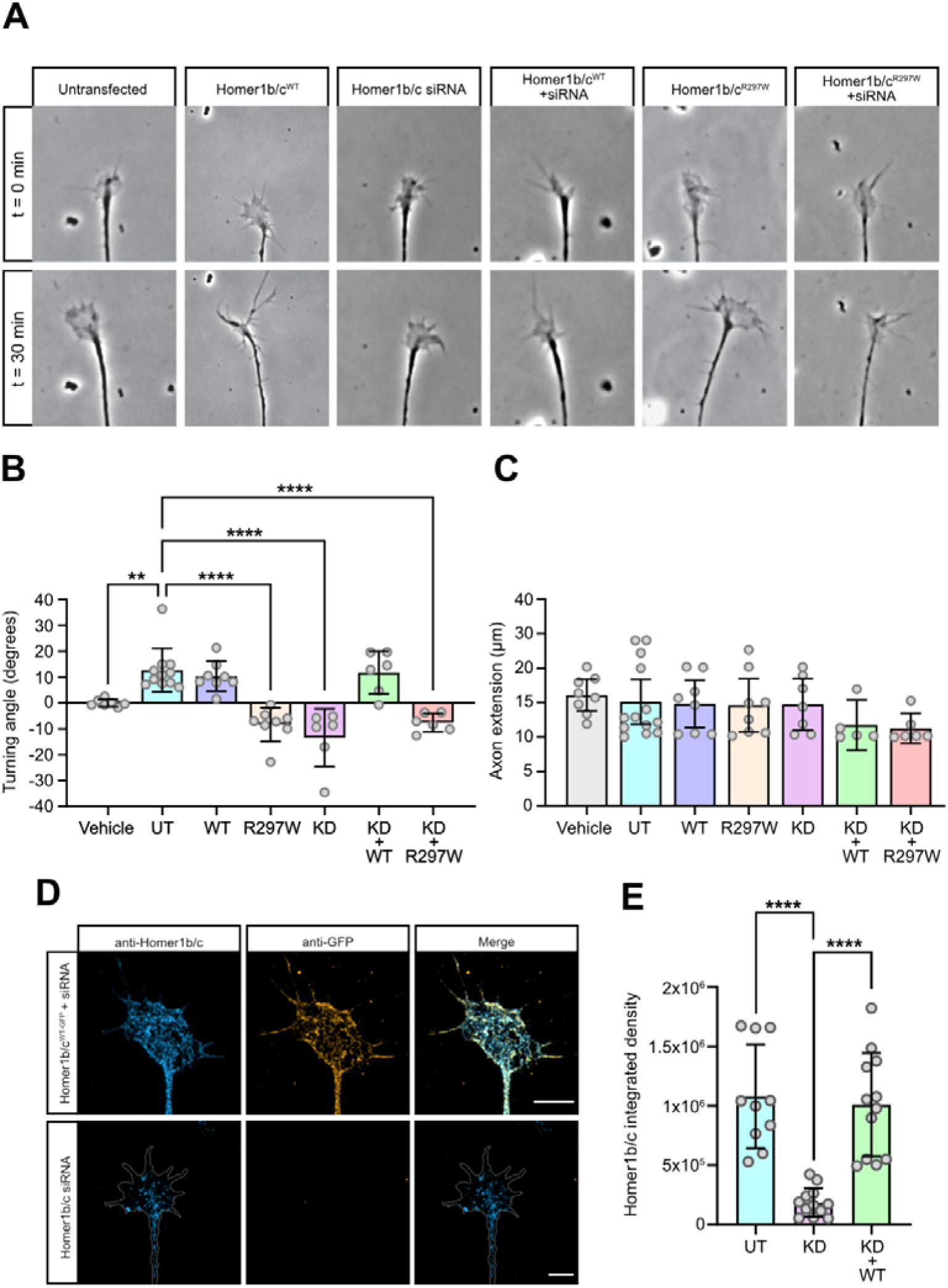
Homer1b/c^R297W^ impairs growth cone turning responses to BDNF. **(A)** Representative time-lapse images of DRG growth cones at start (0 min) and end (30 min) of BDNF-directed turning assay across all groups. Scale bar = 10 µm. **(B)** Quantification of growth cone turning angles elicited by BDNF. Positive angles denote attractive turning and negative angles repulsive turning relative to the source of the BDNF guidance cue. Each data point represents one growth cone, n = 52 cells, >3 experiments/condition, error bars +/- SD (** p < 0.005, **** p < 0.00005, one-way ANOVA with Dunnett’s multiple comparisons test against UT). **(C)** Quantification of mean axonal extension during 30 min turning assay. Each data point represents one growth cone, n = 52 cells, >3 experiments/condition, error bars +/- SD (one-way ANOVA with Tukey’s multiple comparisons test, ns for all comparisons, p > 0.05). **(D)** Representative images of anti-Homer1b/c and anti-GFP immunocytochemistry in Homer1b/c siRNA and Homer1b/c^WT-GFP^ + siRNA growth cones. Scale bars = 10µm. **(E)** Quantification of integrated density of fluorescence of anti-Homer1b/c signal within untransfected, KD and KD+WT growth cones. Each data point represents one growth cone, n = 34 cells, 3 experiments/condition, error bars +/- SD (**** p < 0.0005, one -way ANOVA with Tukey’s multiple comparisons test).

Exogenous expression of Homer1b/c^WT^ alone did not significantly alter growth cone turning relative to the source of a BDNF cue gradient (10.35°) as compared to un-transfected controls (12.67°, p = 0.9669), suggesting that axon guidance was not impaired by overexpression of the protein (Fig. 1B) However, exogenous expression of Homer1b/c^R297W^ in growth cones caused them to be repelled by the normally attractive BDNF gradient (-8.343°, p < 0.0001) (Fig. 1B). In agreement with previous investigations, we observed that endogenous Homer1b/c was required for appropriate guidance towards BDNF, as siRNA-induced knockdown of endogenous

Homer1b/c induced aberrant repulsive turns to a BDNF gradient (-13.42°, p < 0.0001) (Fig. 1B). A vehicle control replacing BDNF with culture medium yielded neither attractive nor repulsive turning responses, confirming that the responses observed in other conditions were specific to BDNF (0.01086°, p = 0.00035) (Fig. 1B). We additionally confirmed that rates of axon extension were not significantly altered within any of the groups, demonstrating that the presence of endogenous Homer1b/c, and exogenous expression of either Homer1b/c^WT^ or Homer1b/c^R297W^ affected only the turning response of growth cones to BDNF, not the rate of axonal extension (Fig. 1C).

We next sought to characterise the effect of exogenous Homer1b/c^WT^ or Homer1b/c^R297W^ expression in the absence of endogenous Homer1b/c. This was accomplished using a combined transfection protocol of overexpression with siRNA targeting endogenous Homer1b/c siRNA with simultaneously exogenous expression of the WT or R297W isoforms. We confirmed successful Homer1b/c knockdown with immunocytochemistry in growth cones fixed at the same timepoint post-transfection as neurons used in the turning assay (Fig. 1D). A robust reduction in Homer1b/c expression was observed in growth cones transfected solely with Homer1b/c siRNA (1.87 x 10^5^ integrated density) compared with both untransfected growth cones (1.08 x 10^6^, p < 0.0001) and growth cones where Homer1b/c expression was rescued by exogenous Homer1b/c^WT^ overexpression (1.01 x 10^6^, p < 0.0001) (Fig. 1E).

In Homer1b/c^siRNA/WT^ growth cones, we found that the appropriate attractive turning response towards the BDNF gradient was rescued by exogenous expression of Homer1b/c^WT^ (11.79°, p = 0.9997) as compared to untransfected controls, demonstrating that its expression is required for guidance to BDNF. By contrast, we found that Homer1b/c^siRNA/R297W^ growth cones continued to be repelled from BDNF (-7.591°, p < 0.0001), suggesting that Homer1b/c^R297W^ was unable to rescue the effect of endogenous Homer1b/c knockdown.

Considered together, these results demonstrate that exogenous expression of Homer1b/c^R297W^ is sufficient to impair growth cone turning responses to BDNF, irrespective of endogenous Homer1b/c expression. Moreover, the induction of aberrantly repulsive motility responses to BDNF by exogenous Homer1b/c^R297W^ expression alone, against even a background of endogenous Homer1b/c, suggests that Homer1b/c^R297W^ may exert a dominant-negative regulatory effect upon normal Homer1b/c function. The failure of Homer1b/c^R297W^ to rescue knockdown-induced aberrant turning responses to BDNF, as compared to successful rescue with Homer1b/c^WT^, strongly suggest that the *HOMER1^R297W^* substitution significantly impairs Homer1b/c^R297W^ function.

### Homer1b/c^R297W^ expression impairs SOCE in sensory neuron growth cones

Growth cone responses to BDNF are SOCE-dependent (Gasperini et al., 2009; Mitchell et al., 2012; Pavez et al., 2019), a process regulated by ligands such as TRPCs, IP_3_Rs, STIM1 and Orai, which contain compatible binding sites for the Homer1 EVH1 domain. We reasoned that the aberrant growth cone turning responses to BDNF gradients observed in sensory neurons expressing Homer1b/c^R297W^ may be caused by an impairment of normal SOCE.

To investigate this, we performed live ratiometric Ca^2+^ imaging using Fura2 AM, and recorded SOCE in growth cones emptied of ER-Ca^2+^ in the presence of Ca^2+^-free extracellular environment and the SERCA inhibitor, cyclopiazonic acid (CPA) (Fig. 2A, B). Growth cones expressing exogenous Homer1b/c^R297W^ exhibited significantly reduced SOCE as compared to UT controls (Homer1b/c^R297W^ Δ0.5626, UT Δ1.040, p = 0.0234). while Homer1b/c^WT^ overexpression had no significant effect upon SOCE compared to UT (Δ0.9624, p = 0.9395) (Fig. 2C). Inhibition of SOCE with 3µM SKF 96365 almost completely abolished the increase in cytosolic Ca^2+^ following restoration of the 2.5mM Ca^2+^ extracellular environment, confirming that the Ca^2+^ flux observed was mediated by SOC channels (Fig. 2C). The amplitude of SOCE was comparable between the Homer1b/c^R297W^ and 3µM SKF 96365 groups, suggesting that SOCE was substantially abrogated by the exogenous expression of Homer1b/c^R297W^.

**Figure 2:**
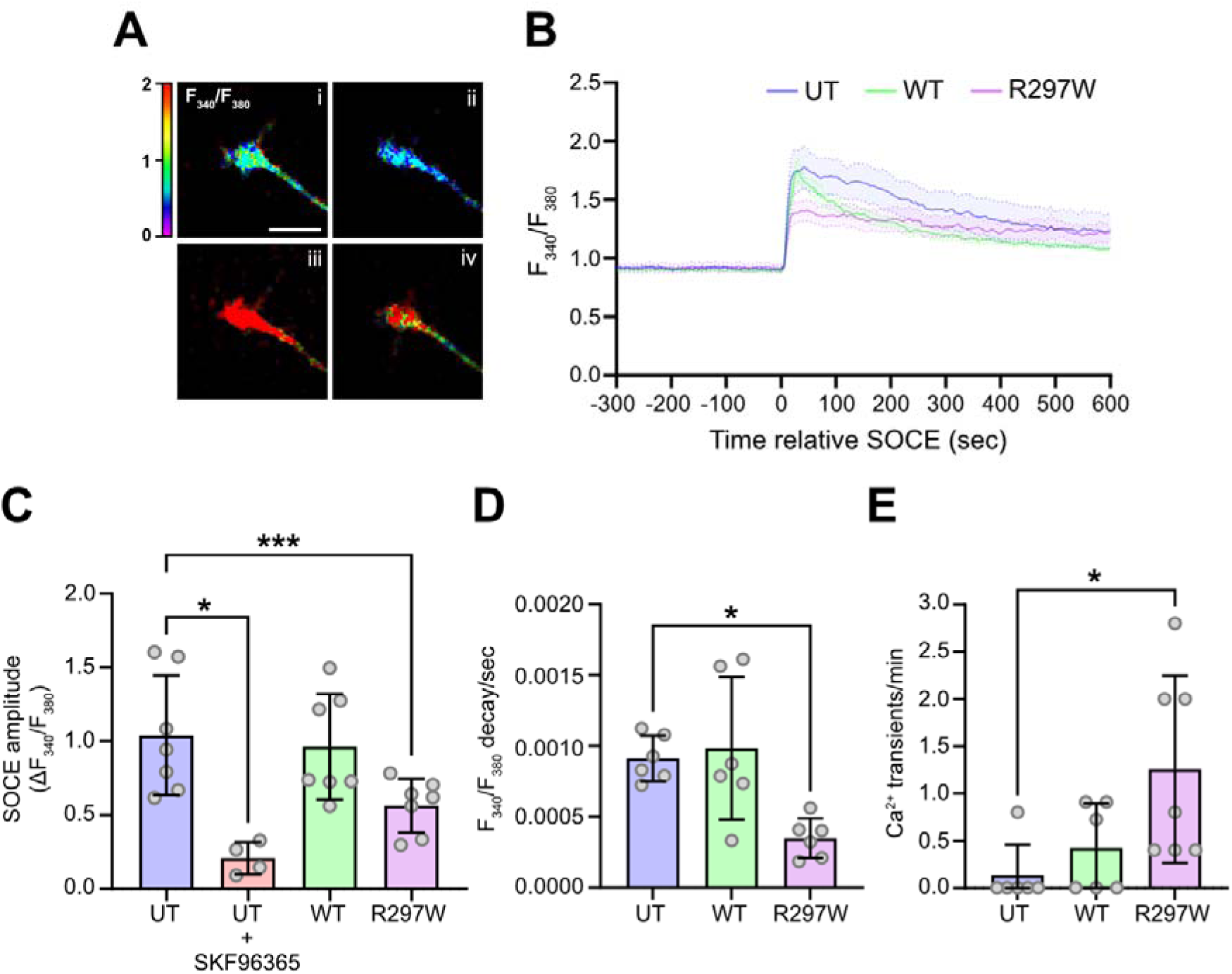
Homer1b/c^R297W^ impairs Ca^2+^ signalling processes in DRG growth cones. **(A)** Representative images of a growth cone during Ca^2+^ imaging at baseline (i), during ER-Ca^2+^ emptying in a Ca^2+^-free extracellular environment in the presence of CPA, during peak SOCE (iii) and on the backslope of the Ca^2+^ increase induced by SOCE. Colour bar denotes range of F_340_/F_380_ ratiometric values on a scale from ∼0 to 2. Scale bar = 10 µm. **(B)** Averaged traces of F_340_/F_380_ levels in all untransfected, Homer1b/c^WT^ and Homer1b/c^R297W^ growth cones to and following SOCE. Traces denote F_340_/F_380_ values recorded at a time interval ranging from -300 seconds before peak SOCE amplitude, and 600 seconds after. Error bars +/- SEM. **(C)** Peak F_340_/F_380_ values during SOCE, calculated as the difference between peak 340/380 following Ca²⁺ re-addition and the final value immediately prior to the re-introduction of a Ca2+ replete environment. Each data point represents one growth cone, n = 25 cells, >3 experiments/condition, error bars +/- SD. (* p < 0.05, *** p < 0.0005, one-way ANOVA with Dunnett’s multiple comparisons test against UT). **(D)** The rate of Ca^2+^ mobilisation per sec following SOCE, calculated based on the rate of F_340_/F_380_ decline from the peak SOCE amplitude to the value at the last frame of recording. Each data point represents one growth cone, n = 20 cells, >3 experiments/condition, error bars +/- SD. (* p < 0.05, one-way ANOVA with Dunnett’s multiple comparisons test against UT). **(E)** The quantity of transient Ca^2+^ increases observed per min during baseline 2.5mM Ca^2+^ replete conditions in WT, UT and R297W growth cones. Each data point represents one growth cone, n = 20 cells, >3 experiments/condition, error bars +/- SD. (* p < 0.05, one-way ANOVA with Dunnett’s multiple comparisons test against UT).

Post-SOCE, cytosolic Ca^2+^ falls as it mobilised to both refill internal Ca^2+^ stores such as the ER via SERCA pumps, and as it is moved to the extracellular environment by ATP-dependent Ca^2+^-channels. Following induction of SOCE, we observed that the rate at which cytosolic Ca^2+^ fell following its peak was reduced in Homer1b/c^R297W^ growth cones (ΔF_340_/F_380_ per/s = 0.0003488) by ∼1/3 compared to both Homer1b/c^WT^ (ΔF_340_/F_380_ per/s = 0.0009840) and UT controls (ΔF_340_/F_380_ per/s = 0.0009122), suggesting that Homer1b/c^R297W^ may impair the functioning of cellular machinery involved in the remobilisation of cytosolic Ca^2+^ following SOCE (Fig. 2D).

Previous work has demonstrated that Homer1b/c is involved in the regulation of spontaneous cytosolic Ca^2+^ transients in growth cones. These transients are sensitive to antagonists of store-operated channels including La^3+^ and SKF-96365, but not voltage-gated calcium channel antagonists, suggesting that the source of the transients is SOC channels (Gasperini et al., 2009). Thus, we were curious as to whether exogenous expression of Homer1b/c^R297W^ may alter spontaneous Ca^2+^ transient activity. We observed a significant increase the frequency of spontaneous Ca^2+^ transients during baseline Ca^2+^ conditions in Homer1b/c^R297W^ growth cones (1.257 transients/min), but not Homer1b/c^WT^ growth cones (0.4242 transients/min), compared to UT controls (0.1333 transients/min) (Fig. 2E). No Ca^2+^ transients were observed in Ca^2+^-free conditions with or without CPA (data not shown), suggesting that these transients represented an extracellular source of Ca^2+^ influx into the cytosol of growth cones.

### Homer1b/c^R297W^ impairs dendritic spine density and SOCE in primary hippocampal neurons

Homer1b/c is required for dendritic spine maturation and stability through its scaffolding of synaptic signalling complexes at synaptic sites. This is antagonised by Homer1a, which competes with Homer1b/c for EVH1-mediated Shank binding and thereby disrupts the greater scaffolding complexes formed through Homer1-Homer1 CC-domain interactions, inhibiting spine density and synaptic activity (Sala et al., 2003; Hayashi et al., 2009). Given that Homer1b/c^R297W^ expression was sufficient to impair Homer1b/c-regulated functions in growth cones, we hypothesised that it may also impair dendritic spine density, plausibly through the impairment of CC-domain mediated tetramerisation of endogenous Homer1b/c. To investigate this, we quantified dendritic spine density in primary hippocampal neurons expressing either transfected Homer1b/c^WT^ or Homer1b/c^R297W^. We observed that Homer1b/c^R297W^ neurons demonstrated significantly lower densities of dendritic spines compared to Homer1b/c^WT^ neurons (1.48/10µm vs 2.31/10µm, p = 0.0179) (Fig. 3A), supporting a model in which CC-domain mediated tetramerisation of Homer1b/c is important for its regulation of dendritic spine density.

**Figure 3:**
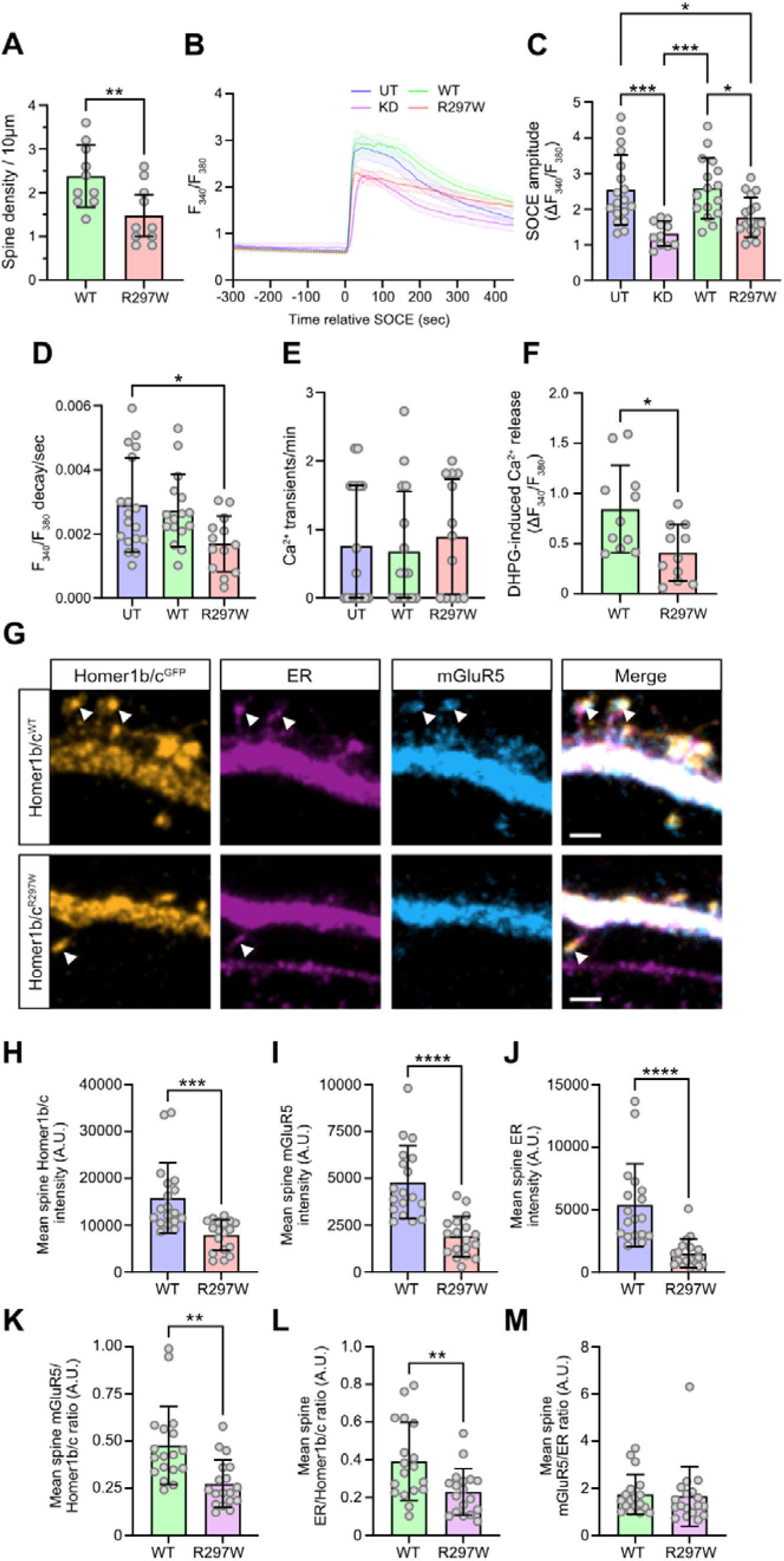
Homer1b/c^R297W^ reduces dendritic spine density, impairs Ca^2+^ signalling and decreases the recruitment of both the ER and mGluR5 to mature hippocampal neuron spines. **(A)** Analysis of dendritic spine density per 10µm in primary hippocampal neurons. Each data point represents one neuron, 10 cells per condition, 3 experiments per condition, error bars +/- SD (** p < 0.005, unpaired t-test). **(B)** Averaged traces of F_340_/F_380_ levels in all untransfected, Homer1b/c^WT^ and Homer1b/c^R297W^ hippocampal neuron somata prior to and following SOCE. Traces denote F_340_/F_380_ values recorded at a time interval ranging from -300 seconds before peak SOCE amplitude, and 450 seconds after. Error bars +/- SEM. **(C)** Peak F_340_/F_380_ values during SOCE, calculated as the difference between peak 340/380 following Ca²⁺ re-addition and the final value immediately prior to the re-introduction of a Ca^2+^ replete environment. Each data point represents one neuron, n = 62 cells, >3 experiments/condition, error bars +/- SD. (* p < 0.05, *** p < 0.0005, one-way ANOVA with Tukey’s multiple comparisons test). **(D)** The rate of Ca2+ mobilisation per sec following SOCE, calculated based on the rate of F_340_/F_380_ decline from the peak SOCE amplitude to the value at the last frame of recording. Each data point represents one neuron, n = 46 cells, 3 experiments/condition, error bars +/- SD. (* p < 0.05, one-way ANOVA with Dunnett’s multiple comparisons test against UT). **(E)** The quantity of transient Ca2+ increases observed per min during baseline 2.5mM Ca2+ replete conditions in WT, UT and R297W hippocampal neurons. Each data point represents one cell, n = 46 cells, 3 experiments/condition, error bars +/- SD. (Krusal-Wallis test with Dunnett’s multiple comparisons test against UT, ns for all comparisons, p > 0.05). **(F)** Peak F_340_/F_380_ values elicited by bath application of 3,5-DHPG, calculated as the difference between baseline prior to addition and the peak 340/380 value post-DHPG application. 11 cells per condition, >3 experiments per condition, errors bars +/- SD (* p < 0.005, unpaired t-test). **(G)** Representative images of Homer1b/c^WT^ and Homer^R297W^ overexpressing hippocampal neurons, immunolabelled for anti-GFP (Homer1b/c^GFP^), anti-mCherry (KDEL-mCherry for ER) and anti-mGluR5. Arrowheads point to spine heads where accumulation of Homer1b/c^GFP^, ER and mGluR5 is evident. Scale bars = 2µm. **(H-J)** Quantification of mean intensity of Homer1b/c^GFP^, mGluR5 and ER in WT and R297W overexpression conditions. Each data point represents dendritic average for all spines analysed, with one dendrite analysed per neuron, n = 18 neurons, 3 experiments per condition, error bars +/- SD (*** p < 0.0005, **** p < 0.00005, unpaired t-test). **(K-L)** Quantification of mGluR5 and ER (integrated density, A.U.) present within dendritic spine heads normalised per unit of Homer1b/c^WT-GFP^. Each data point represents dendritic average for all spines analysed, with one dendrite analysed per neuron, n = 18 neurons, 3 experiments per condition, error bars +/- SD. (** p < 0.005, unpaired t-test). **(M)** Quantification of average mGluR5 expression present within dendritic spine heads, normalised spine head ER (integrated density, A.U.). Each data point represents dendritic average for all spines analysed, with one dendrite analysed per neuron, n = 18 neurons, 3 experiments per condition, error bars +/- SD (unpaired t-test, ns difference in comparison, p > 0.05).

The recruitment of proteins involved in SOCE is a crucial process in regulating Ca^2+^ signalling and hence spine plasticity and development. Both STIM1 and Orai1 function is necessary for the formation of new dendritic spines and for the shaping of spine maturation during early spinogenesis (Korkotian et al., 2017; Kushnireva et al., 2021). Having found evidence of reduced spine density in Homer1b/c^R297W^-expressing neurons and having observed impaired SOCE in sensory neuron growth cones, we hypothesised that SOCE functioning may also be impaired in hippocampal neurons, potentially contributing to the decreased spine density induced by Homer1b/c^R297W^. To investigate this, we applied our established SOCE protocol to conduct live Ca^2+^ imaging in primary hippocampal neurons (Fig. 3B).

Homer1b/c^R297W^ expression significantly decreased the amplitude of SOCE recorded in hippocampal neuron somata (ΔF_340_/F_380_ = 1.744,) compared to both untransfected controls (ΔF_340_/F_380_ = 2.540, p = 0.0210) and neurons overexpressing Homer1b/c^WT^ (ΔF_340_/F_380_ = 2.587, p = 0.0165). In contrast, overexpression of Homer1b/c^WT^ did not significantly alter average SOCE amplitudes compared to UT controls (p = 0.9979) (Fig. 3C). We next asked whether reducing endogenous Homer1b/c expression would similarly attenuate SOCE amplitude as with exogenous Homer1b/c^R297W^ neurons. Knockdown of endogenous Homer1b/c with Homer1b/c siRNA significantly inhibited SOCE (ΔF_340_/F_380_ = 1.317) compared to both untransfected controls (p = 0.0005) and Homer1b/c^WT^ neurons (p = 0.0004) but produced comparable SOCE to that observed in Homer1b/c^R297W^ conditions (p = 0.4101) (Fig. 3C).

In agreement with our earlier observations in growth cones, exogenous expression of Homer1b/c^R297W^ (ΔF_340_/F_380_ per/s = 0.001695) also significantly reduced the rate of Ca^2+^ remobilisation following SOCE compared to UT conditions (ΔF_340_/F_380_ per/s = 0.002905, p = 0.0017), whereas exogenous Homer1b/c^WT^ expression induced no changes (ΔF_340_/F_380_ per/s = 0.002732, p = 0.8864) (Fig. 3D).

In combination with the SOCE experiments conducted in sensory neuron growth cones, these results confirm that functional Homer1b/c is necessary for SOCE. Moreover, this data suggests that the integrity of the CC domain of Homer1b/c is required for the functioning of Homer1b/c in SOCE, and that expression of Homer1b/c^R297W^ is sufficient to impair the likely scaffolding of constituent SOCE proteins by endogenous Homer1b/c, as evidenced by the similar reduction in SOCE observed between knockdown conditions and those where Homer1b/c^R297W^ is exogenously expressed.

### Homer1b/c^R297W^ impairs both group-I mGluR-mediated Ca^2+^ signalling and the recruitment of both the ER and mGluR5 to mature hippocampal neuron spines

Having found that Homer1b/c^R297W^ impaired SOCE in hippocampal neurons, we next asked whether this impairment extended to other Ca^2+^ signalling processes involved in synaptic plasticity and the regulation of dendritic spines. Homer1b/c links group-I mGluR activation to downstream intracellular Ca^2+^ release, a complex process involving ER-Ca^2+^ release via IP3Rs, extracellular Ca^2+^ influx via VGCCs and TRPCs, and further Ca^2+^ influx via SOCs following ER-Ca^2+^ mobilisation. (Tu et al., 1998; Bianchi et al., 1999; Gee et al., 2003; González-Sánchez et al., 2017). Notably, the scaffolding complexes formed by Homer1b/c with mGluRs and other PSD proteins are necessary for proper regulation of group-I mGluR-mediated Ca^2+^ signalling. We therefore hypothesised that exogenous Homer1b/c^R297W^ expression would impair group-I mGluR-mediated Ca^2+^ release, due to its disruption of endogenous Homer1b/c scaffolding complexes.

Group-I mGluRs were activated by bath application of 200µM of 3,5-DHPG and the resulting rise in cytosolic Ca^2+^ was recorded in hippocampal dendrites using ratiometric F_340_/F_380_ imaging. The temporal dimensions of 3,5-DHPG-induced increases were similar between neurons expressing either Homer1b/c^WT^ or Homer1b/c^R297^, with cytosolic Ca^2+^ increasing above baseline levels for ∼30s following application of 3,5-DHPG (data not shown), but the amplitude of Ca^2+^ rises post mGluR-activation were significantly blunted in neurons expressing Homer1b/c^R297W^ (ΔF_340_/F_380_ = 0.356) than those observed in Homer1b/c^WT^ neurons (ΔF_340_/F_380_ = 0.722, p = 0.0158) (Fig. 3F). Given that Homer1a overexpression reduces DHPG-mediated Ca^2+^-release by disrupting the scaffolds coupling mGluRs to ER-bound IP_3_Rs via Homer1b/c (Kammermeier and Worley, 2007), the dampened Ca^2+^ response to 3,5-DHPG suggests that the Homer1b/c-mGluR-IP3R signalling complex could be similarly disrupted by expression of Homer1b/c^R297W^.

Homer1b/c, in combination with Shank, facilitates the formation of molecular scaffolding complexes which recruit ER cisternae to dendritic spines, whereas loss of Shank, Homer1b/c, or increasing expression of Homer1a inhibits ER recruitment (Sala et al., 2001, 2003, 2005). Reduced ER recruitment is in turn associated with reductions in both the density and size of dendritic spines, alongside surface expression of NMDA and AMPA receptors. We therefore hypothesised that Homer1bc^R297W^ expression would reduce ER recruitment to spines via disassembly of these scaffolding complexes. Furthermore, our observation of impaired Ca^2+^-responses elicited by 3,5-DHPG suggested that mGluR-ER scaffolds may be disrupted by the expression of Homer1b/c^R297W^. Given that long term depression (LTD) induced by mGluR-activation has been reported to be restricted to mature, ER-containing spines (Holbro et al., 2009; Perez-Alvarez et al., 2020), we hypothesised that group-I mGluR would be reduced within spines where Homer1b/c^R297W^ was present.

To test this, mouse primary hippocampal neurons were co-transfected with KDEL-mCherry to visualise ER cisternae and either Homer1b/c^WT-GFP^ or Homer1b/c^R297W-GFP,^ then immunolabelled with anti-GFP, anti-mCherry and anti-mGluR5 antibodies (Fig. 3G). After correcting for differences in spine volume, we found that Homer1b/c^WT-GFP^ containing spines demonstrated significantly greater Homer1b/c^GFP^ expression (15845 A.U., p = 0.0002), ER infiltration (5377 A.U., p < 0.0001) and mGluR5 expression (4790 A.U., p = 0.0165) than Homer1b/c^R297W-GFP^ containing spines (Homer1b/c^GFP^ = 7910 A.U., ER = 1519 A.U, mGluR5 = 1895 A.U.) (Figs. 3H-J).

Because the recruitment of ER to dendritic spines has previously been positively correlated with Homer1b/c expression levels (Sala et al., 2001), we next sought to confirm that the decreased spine ER and mGluR5 expression within the R297W group was not attributable to the lower relative Homer1b/c expression within Homer1b/c^R297W-GFP^ spines. For each spine head, we calculated ratios of integrated density for mGluR5 intensity per unit of Homer1b/c intensity, as well as ER/Homer1b/c and mGluR5/ER. Following this normalisation, Homer1b/c^WT-GFP^ spines still retained significantly more ER (0.3916 ER/Homer1b/c, p = 0.0072) and mGluR5 (0.4770 mGluR5/Homer1b/c, p = 0.0011) per unit of Homer1b/c^GFP^ than Homer1b/c^R297W-GFP^ spines (ER/Homer1b/c = 0.2299, mGluR5/Homer1b/c = 0.2739) (Figs. 3K-L). The mGluR5/ER ratio did not differ between the genotypes (Fig. 3M).

Taken together, these results show that Homer1b/c^R297W^ expression reduces both mGluR5 expression and ER recruitment at dendritic spines. Since both Homer1b/c tetramerisation and a functional CC domain are required for efficient synaptic targeting (Hayashi et al., 2006), the reduced expression of Homer1b/c^R297W-GFP^ within spines suggests that the CC domain function – and by extension its capacity for tetramerisation – is impaired relative to Homer1b/c^WT^, alongside its capacity to recruit both mGluR5 and the ER to spines.

### Homer1b/c^R297W^ disrupts nanoscale coupling to mGluR5 and STIM1 in growth cones

The balance of Homer1a and Homer1b/c binding to group-I mGluRs is among the most thoroughly studied of all Homer1 functions, having been described in the regulation of activity dependent plasticity, metaplasticity and homeostatic scaling, chronic stress, traumatic brain injury, sleep and network hyperexcitability, and many other domains (Brakeman et al., 1997; Tu et al., 1998; Xiao et al., 1998; Kammermeier et al., 2000; Mao et al., 2005; Kammermeier and Worley, 2007; Ronesi et al., 2012; Luo et al., 2014; Guo et al., 2016; Aloisi et al., 2017; Buscemi et al., 2017; Diering et al., 2017; Hu et al., 2017; Bockaert et al., 2021; Guerrero and Turrigiano, 2025b). Having found evidence that Homer1b/c^R297W^ downregulated both Ca^2+^ flux following group-I mGluR activation and mGluR5 recruitment to dendritic spines, we hypothesised that these functional alterations could be correlated with a direct uncoupling of Homer1b/c^R297W^ from mGluR5. Given that the tetramerisation of Homer1b/c has been shown previously to be necessary for its co-clustering with group-I mGluRs (Hayashi et al., 2006), we reasoned that evidence of significant uncoupling of Homer1b/c^R297W^ from mGluR5 would suggest an impairment in capacity for Homer1b/c^R297W^ to properly form homotetramers.

To investigate this, sensory neurons growth cones were transfected with Homer1b/c^WT-GFP^ or Homer1b/c^R297W-GFP^ and immunolabelled with anti-GFP and anti-mGluR5 antibodies, while untransfected growth cones were labelled anti-Homer1b/c. We elected to return to sensory neuron growth cones to study the colocalisation of Homer1b/c and mGluR5 owing to the exceptional optical properties due to their thin and flat structure. We used statistical object distance analysis (SODA) (Lagache et al., 2018) to compute a coupling probability (CP) for all untagged or untagged Homer1b/c with mGluR5 puncta within the growth cone. Homer1b/c^R297W-GFP^ puncta were significantly less likely to be coupled with mGluR5 (CP = 0.3593) than were both Homer1b/c^WT-GFP^ (CP = 0.5437, p = 0.0003) and untransfected Homer1b/c (CP = 0.5332, p = 0.0007).

Although diffraction-limited imaging revealed a clear spatial uncoupling of Homer1b/c^R297W-GFP^ and mGluR5, the level of resolution offered by this modality cannot resolve whether this reflects a fundamental reorganisation of their nanoscale spatial relationship. To address this limitation, we used direct stochastic optical reconstruction microscopy (dSTORM) to further probe with super-resolution the distribution of Homer1b/c relative to known epitopes at the level of individual protein assemblies. DRG growth cones were transfected with Homer1b/c^WT-GFP^ or Homer1b/c^R297W-GFP^ and immunolabelled with anti-GFP, along with either an anti-STIM1 or anti-mGluR5. Resolved dSTORM images revealed that Homer1b/c, STIM1, and mGluR5 were widely distributed throughout the growth cone, consistent with our earlier findings (Fig. 4C).

**Figure 4:**
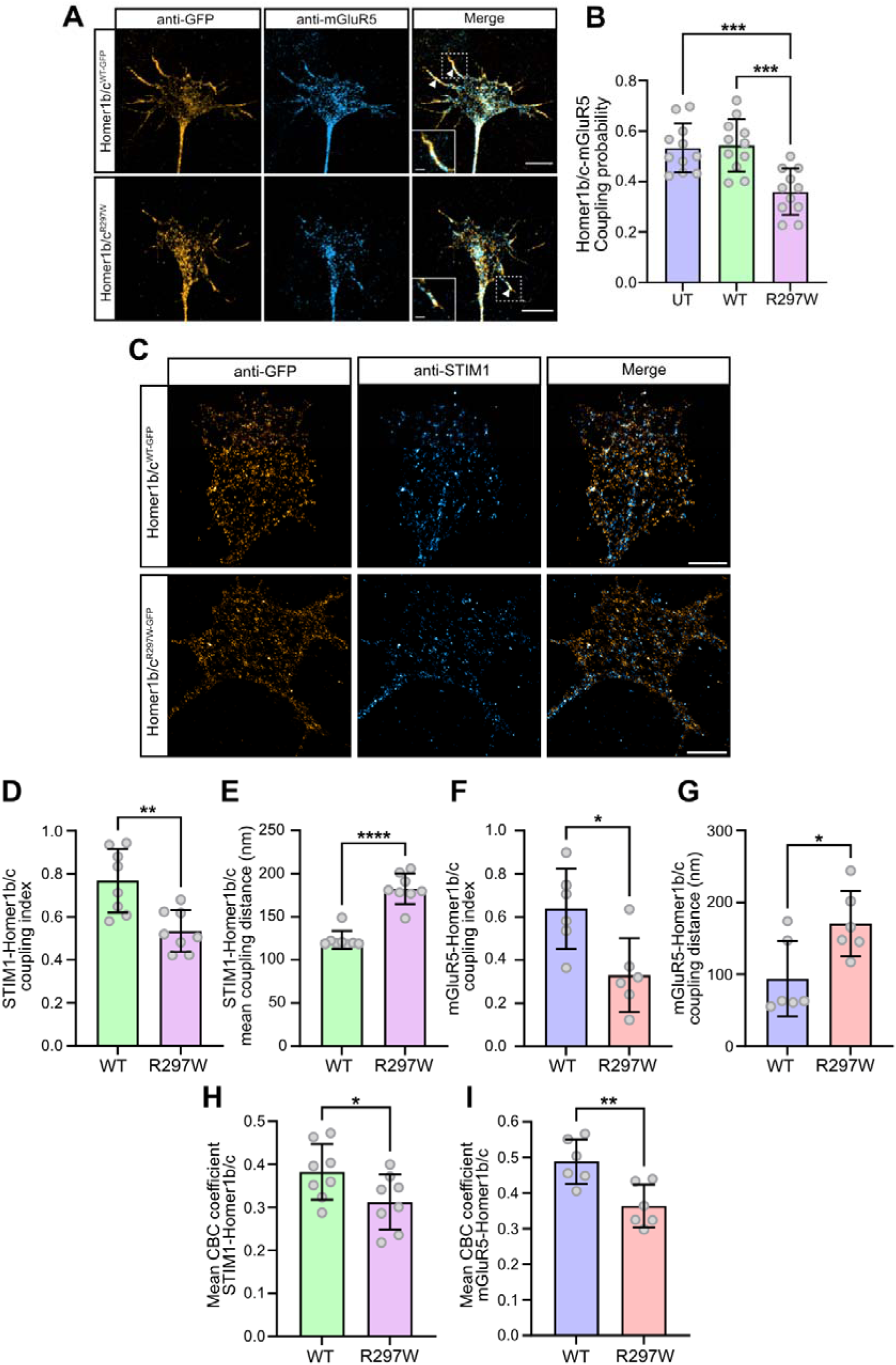
Homer1b/c^R297W^ is de-coupled from STIM1 and mGluR5 in sensory neuron growth cones. **(A)** Representative images of DRG sensory neuron growth cones immunolabelled with an anti-mGluR5 antibody and anti-GFP targeting transfected Homer1b/c^WT-GFP^ or Homer1b/c^R297W-GFP^. Arrowheads point to regions of colocalised Homer1b/c^GFP^ and mGluR5 puncta within growth cone filopodia. Insert boxes are expanded views of filopodia marked by dashed square in merged images. Scale bars = 10µm. **(B)** Quantification of computed coupling probability between Homer1b/c (UT) or Homer1b/c^GFP^ (WT and R297W overexpression) and mGluR5 puncta within sensory neuron growth cones. Each data point represents one growth cone, n = 11 per condition, 3 experiments per condition, error bars +/- SD (*** p < 0.0005, one-way ANOVA with Tukey’s multiple comparisons test). **(C)** Representative reconstructed dSTORM image of Homer1b/c^WT-GFP^ and Homer1b/c^R297W-GFP^ growth cones immunolabelled with anti-GFP and anti-STIM1. Scale bars = 2μm. **(D)** Quantification of mean SODA computed coupling indices describing the association strength between STIM1 and Homer1b/c^GFP^ puncta in Homer1b/c^WT^ and Homer1b/c^R297W^ expressing growth cones in resolved dSTORM images. Each data point represents one growth cone, n = 8 cells per condition, 3 experiments/condition, error bars +/- SD (** p < 0.005, unpaired t-test with Welsch’s correction). **(E)** Mean distance in nanometres between spatially coupled STIM1 and Homer1b/c^GFP^ in Homer1b/cWT and Homer1b/cR297W expressing growth cones in resolved dSTORM images. Each data point represents one growth cone, n = 8 cells per condition, 3 experiments/condition, error bars +/- SD (**** p < 0.00005, unpaired t-test with Welsch’s correction). **(F)** Quantification of mean SODA computed coupling indices describing the association strength between mGluR5 and Homer1b/c^GFP^ puncta in Homer1b/c^WT^ and Homer1b/c^R297W^ expressing growth cones in resolved dSTORM images. Each data point represents one growth cone, n = 6 cells per condition, 3 experiments/condition, error bars +/- SD (* p < 0.05, unpaired t-test with Welsch’s correction). **(G)** Mean distance in nanometres between spatially coupled STIM1 and Homer1b/c^GFP^ in Homer1b/c^WT^ and Homer1b/c^R297W^ expressing growth cones in resolved dSTORM images. Each data point represents one growth cone, n = 6 cells per condition, 3 experiments/condition, error bars +/- SD (* p < 0.05, unpaired t-test with Welsch’s correction). **(H)** Quantification of mean STIM1-Homer1b/c^GFP^ CBC coefficients per cell in WT and R297W growth cones. Each data point represents one growth cone, n = 8 cells per condition, 3 experiments per condition, error bars +/- SD (* p < 0.05, unpaired t-test with Welsch’s correction). **(I)** Quantification of mean mmGluR5-Homer1b/cGFP CBC coefficients per cell in WT and R297W growth cones. Each data point represents one growth cone, n = 6 cells per condition, 3 experiments per condition, error bars +/- SD (** p < 0.05, unpaired t-test with Welsch’s correction).

We utilised 2D-SODA STORM (Lagache et al., 2018) to analyse dSTORM localisation data and calculate coupling indices (CI), the probability weighted association strength between STIM1 or mGluR5 localisations with Homer1b/c^GFP^ localisations within a given growth cone. We found that the mean STIM1-Homer1b/c CI was significantly increased between STIM1 and Homer1b/c^WT-GFP^ localisations (CI 0.7675) as compared to STIM1-Homer1b/c^R297W-GFP^ localisation (0.5314, p = 0.0029) (Fig. 4D). To further probe the nanoscale localisation of these proteins, we also sought to compare the mean distance in nanometres between significantly coupled STIM1 and Homer1b/c^GFP^ puncta. There was a significant increase in the mean coupling distance between spatially associated STIM1 and Homer1b/c^R297W-GFP^ localisations (182.7nm) as compared to the distance separating spatially coupled STIM1 and Homer1b/c^WT-GFP^ localisations (123.4nm, p < 0.0001) (Fig 4. E). Similar differences between WT and R297W groups were observed in the case of mGluR5-Homer1b/c^GFP^ coupling. The average association strength of spatially coupled of mGluR5-Homer1b/c^WT-GFP^ localisations were significantly elevated (CI = 0.6387) as compared to mGluR5-Homer1b/c^R297W-GFP^ (CI = 0.33806, p = 0.0138) (Fig. 4F), and a significantly increased distance in nm between spatially coupled Homer1b/c^R297W-GFP^ and mGluR5 localisations was observed (170.74nm), as compared to that calculated between significantly coupled mGluR5 and Homer1b/c^GFP^ localisations (93.94nm, p = 0.0228) (Fig. 4G).

While SODA has been applied elsewhere to quantitatively assess protein coupling within synaptic compartments, we are not aware of its prior application with neuronal growth cones. We therefore complemented SODA with additional coordinate-based colocalisation analysis (CBC) to provide another independent measure of coupling strength for comparison.

For each detected STIM1 or mGluR5 localisation, a CBC coefficient ranging from -1 to 1 was calculated relative to Homer1b/c^GFP^, where higher positive values represented stronger spatial association and localisation between STIM1/mGluR5 and Homer1b/c. To ease comparisons between growth cones with substantially differing numbers of detected STIM1/mGluR5 localisations, we expressed the colocalisation coefficient as the mean CBC coefficient per cell, calculated as the frequency-weighted average across the -1 to 1 CBC distribution. We compared the mean CBC coefficient per cell for values filtered between 0.15 and 1 to focus on colocalised STIM1/mGluR5 and Homer1b/cR297W^GFP^ puncta in Homer1b/c^WT^ and Homer1b/c^R297W^ growth cones. Both the mean STIM1-Homer1b/c^R297W-GFP^ and mGluR5-Homer1b/c^R297W-GFP^ CBC coefficients per cell were significantly decreased (CBCs = 0.3124 and 0.3638, respectively) compared to the strength of colocalisation between the two epitopes and Homer1b/c^WT-GFP^ (STIM1 CBC = 0.3822, p = 0.0477, mGluR5 CBC = 0.4882, p = 0.0057) (Figs. 3H, 3I).

The application of SODA coupling analysis and Voronoi tessellation-based clustering to neuronal growth cones represents, to our knowledge, a novel analytical approach for the interrogation of protein-protein interactions in this neuronal compartment, which we have used here to investigate nanoscale-level protein coupling and colocalisation. Considered together, both our diffraction-limited and super-resolution enabled analyses report that the Homer1b/c^R297W^ variant demonstrates significantly impaired coupling between it and both mGluR5 and STIM1 in sensory neuron growth cones.

### Homer1b/c^R297W^ impairs the nanoscale clustering of protein scaffolds within dendritic spine heads

Appropriate scaffolding of signalling proteins is a prerequisite for their assembly into functional nanodomains within restricted cellular compartments, including dendritic spines. Having observed that Homer1b/c^R297W^ exhibits impaired coupling to key ligands involved in Ca^2+^ signalling within growth cones, we next asked whether this scaffolding deficit extended to an alteration in the local clustering of Ca^2+^ signalling components within dendritic spine heads. Because Ca^2+^ signalling within mature spine is supported by ER-associated signalling complexes, we focused on the clustering of mGluR5 together with IP_3_R and STIM2, with mediate ER-Ca^2+^ release and sustained SOCE at spines, respectively.

Hippocampal neurons were transfected with Homer1b/c^WT-GFP^ or Homer1b/c^R297W-GFP^ and immunolabelled with an anti-GFP antibody in addition to anti-STIM2, anti-IP_3_R or anti-mGluR5. Clusters of mGluR5, STIM2 and IP_3_R within dendritic spine heads were classified with Voronoi segmentation from spectrally de-mixed dSTORM data and compared between Homer1b/c^WT-GFP^ and Homer1b/c^R297W-GFP^ neurons (Fig. 5A). Because clusters of mGluR5, STIM2 and IP3R clusters were considerably more frequently within Homer1b/c^WT-GFP^ spines than Homer1b/c^R297W-GFP^ spines, we elected to average the data describing clusters across all spines that were analysed for a given cell. Expression of Homer1b/c^R297W^ was associated with a marked reduction in the density of clusters of each Ca^2+^ signalling protein within dendritic spine heads. Compared to Homer1b/c^WT-GFP^ spines, Homer1b/c^R297W^ spines exhibited significantly lower densities of mGluR5 (283.5 loc/µm^2^ vs 402 loc/µm^2^, p = 0.0284) (Fig. 5B), STIM2 (468.6 loc/µm^2^ vs 887.4 loc/µm^2^, p = 0.01) (Fig. 5C), and IP_3_R (539.1 loc/µm^2^ vs 1679 loc/µm^2^, p = 0.0036) (Fig. 5D). In contrast, the mean number of localisations per cluster did not significantly differ between Homer1b/c^WT^ and Homer1b/c^R297W^ spines for mGluR5 (Fig. 5E), IP3R (Fig. 5F) or STIM2 (Fig. 5G), which suggested that while the overall number of protein localisations did not change between the conditions, the nanoscale distribution of these proteins is impaired.

**Figure 5:**
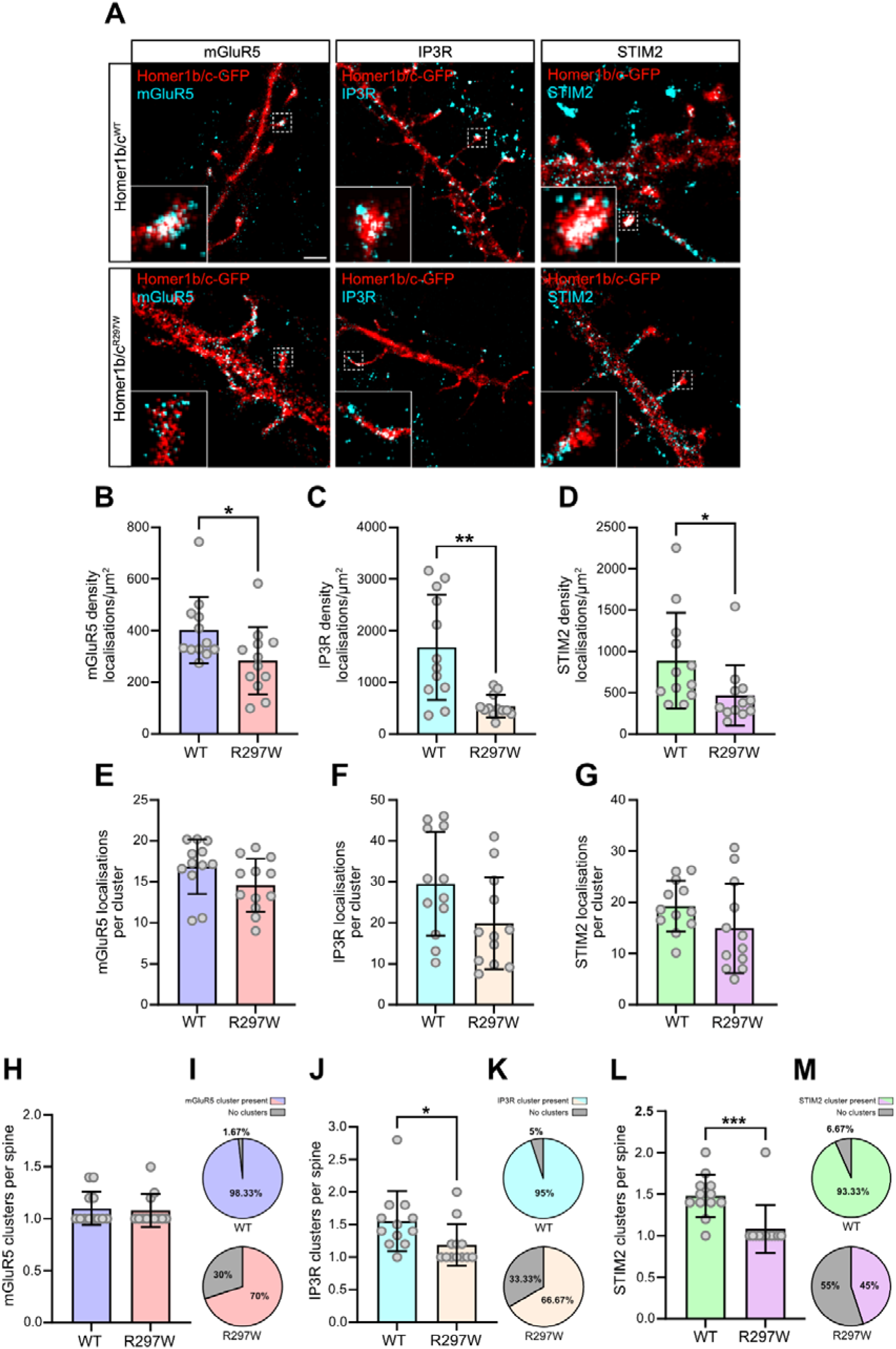
The density of mGluR5, STIM2 and IP3R clusters is reduced within spine heads containing Homer1b/c^R297W^. **(A)** Representative images reconstructed from dSTORM of Homer1b/c^WT-GFP^ and Homer1b/c^R297W-GFP^ neurons immunolabelled with anti-GFP and either mGluR5, IP3R or STIM2. Images shown represent 15mm pixel size reconstructions from processed and demixed acquisitions. Inserts show magnified views of Homer1b/c^GFP^ and target epitopes in spine heads, sourced from the boxed regions in the original view. Scale bar = 2µm. **(B)** The density of mGluR5 clustering in Homer1b/c^WT^ and Homer1b/c^R297W^ spine heads, as quantified by the number of mGluR5 localisations resolved per µm^2^ of each total cluster size defined by Voronoi segmentation analysis. Each data point represents the average of 5 spines per cell, n = 12 per condition, 3 experiments/condition, error bars +/- SD (* p < 0.05, Mann-Whitney U-test). **(C)** The density of IP3R clusters in Homer1b/c^WT^ and Homer1b/c^R297W^ spines, also quantified by the number of IP3R localisations resolved per µm^2^ of each total cluster size. Each data point represents the average of 5 spines per cell, n = 12 per condition, 3 experiments/condition, error bars +/- SD (** p < 0.005, Mann-Whitney U-test) **(D)** Quantified clustering of STIM2 in Homer1b/c^WT^ and Homer1b/c^R297W^ spines, defined as the number of STIM2 localisations resolved per µm^2^ of each total cluster size defined by Voronoi analysis. Each data point represents the average of 5 spines per cell, n = 12 per condition, 3 experiments/condition, error bars +/- SD (*

We next quantified both the proportion of dendritic spines containing detectable clusters of each target protein, as well as the mean number of clusters that could be identified per spine. STIM2 clusters were detected in the majority of Homer1b/c^WT-GFP^ spines (93.33%), but in fewer than half of Homer1b/c^R297W-GFP^ spines (45%) (Fig. 5I). A similar pattern was also observed for IP3R, where clusters were present in 95% of Homer1b/c^WT-GFP^ spines, compared with 66.67% of Homer1b/c^R297W-GFP^ spines (Fig. 5K). In the case of mGluR5 clusters, we also observed a reduced proportion of spines that contained at least one mGluR5 cluster when comparing Homer1b/c^R297W^ and Homer1b/c^WT^ neurons (70% vs 98.33%) (Fig. 5M). Additionally, among spine heads that contained at least one STIM2 cluster, Homer1b/c^WT-GFP^ spines contained significantly more clusters per spine on average as compared to Homer1b/c^R297W-GFP^ spines (1.479 vs 1.083, p = 0.0003) (Fig. 5E). The mean number of IP_3_R clusters per spine was also significantly reduced in in the R297W neurons (1.189 vs 1.553, p = 0.0106) (Fig 5F). By contrast, the mean number of mGluR5 clusters per spine did not differ between the groups (1.079 vs 1.100, p = 0.8202) (Fig. H), suggesting that, while we observed mGluR5 to be less commonly expressed in spine heads, when it was successfully recruited, it was shepherded into a number of clusters comparable between Homer1b/c^WT^ and Homer1b/c^R297W^ spines. Such an effect could reflect a state of increased mGluR5 mobility, as occurs when Homer1b/c mGluR5 scaffolds are disrupted (Aloisi et al., 2017), and may suggest the presence of intramolecular mechanisms involved in clustering and scaffolding of mGluR5 that are distinct from those with which IP3R and STIM2 associate.

Collectively, these results demonstrate that Homer1b/c^R297W^ disrupts the recruitment and nanoscale organisation of key proteins involved in facilitating Ca^2+^ signalling within the dendritic spine head. That the density of mGluR5, STIM2 and IP3R clusters was decreased with the expression of Homer1b/c^R297W^ in spite of no significant change to the overall number of localisations representing those proteins strongly suggests that clusters of mGluR5/STIM2/IP3R are significantly more compact within Homer1b/c^WT^ spines than Homer1b/c^R297W^ spines. This finding has important implications for the functioning of these proteins, as the clustering of glutamatergic receptors and their Ca^2+^ signalling effectors into nanodomains contributes to the efficiency of their functioning.

## Discussion

In this study, we have functionally characterised a previously unreported de novo human *HOMER1* variant, *HOMER1^R297W^*, and demonstrate that its expression is associated with a broad disruption of Homer1b/c-dependent processes. Our finds support a model in which the R297W substitution within the coiled-coil domain impairs Homer1b/c tetramerisation, disrupting the formation of the scaffolding complexes necessary for the spatial organisation of Ca^2+^ signalling machinery within neurons. Super resolution imaging revealed that Homer1b/c^R297W^ coupling to mGluR5 and STIM1 is reduced in growth cones, and that the clustering of mGluR5, STIM2 and IP3R within dendritic spine heads is diminished, consistent with a loss of scaffold-dependent compartmentalisation. These scaffolding deficits were accompanied by attenuated SOCE in both sensory neuron growth cones and hippocampal neurons, blunted Ca^2+^ responses to group-I mGluR activation, and reduced recruitment of ER and mGluR5 to mature spines. At the cellular level, Homer1b/c^R297W^ expression reduced dendritic spine density and converted normally attractive axon guidance responses to repulsion – phenotypes consistent with the downstream consequences of impaired Ca^2+^ regulation during neuronal development. The convergence of these phenotypes points to a requirement for intact tetrameric Homer1b/c scaffolding complexes in the regulation of neuronal Ca^2+^ homeostasis across both pre- and postsynaptic compartments.

### Coiled-coil domain integrity and the scaffolding requirement for Homer1b/c function

Our work here demonstrates that a single substitution within the coiled-coil domain is sufficient to broadly impair Homer1b/c function in a manner consistent with defective tetramerisation. The R297 residue mediates intermolecular salt bridges within the C1 dimeric interface of the Homer1b/c coiled-coil (Hayashi et al., 2009), and its replacement with an uncharged tryptophan is predicted to destabilise these contacts. While we did not directly measure the multimerisation state of Homer1b/c^R297W^, several lines of evidence support impaired tetramer formation. Homer1b/c^R297W^ was significantly less coupled with mGluR5 in growth cones, consistent with the established requirement of Homer1b/c tetramerisation for mGluR co-clustering (Hayashi et al., 2006). The reduced density of mGluR5, STIM2 and IP3R clusters within Homer1b/c^R297W^ spine heads, despite unchanged numbers of localisations per cluster, further suggests that these proteins are less efficiently compartmentalised into the compact signalling domains characteristic of intact Homer1b/c scaffolds. Critically, Homer1b/c^R297W^ impaired both axon guidance and SOCE against a background of endogenous wild-type protein, whereas additional Homer1b/c^WT^ expression had no effect. This pattern – wherein Homer1b/c^R297W^ disrupts endogenous function while Homer1b/c^WT^ overexpression is tolerated – is consistent with a dominant-negative mechanism of action and argues against a purely loss-of-function interpretation. We posit a model under which Homer1b/c^R297W^ may incorporate into heteromeric complexes with endogenous Homer1b/c, reducing the proportion of functional tetramers available for scaffold assembly, analogous to the behaviour of the engineered dimeric variant Homer1b/c^I332R/I337E^ (Hayashi et al., 2009).

The impaired axon guidance, attenuated SOCE, blunted mGluR-dependent Ca²⁺ signalling, decreased spine density and reduced synaptic targeting of mGluR5 caused by Homer1b/c^R297W^ expression parallel those reported following either Homer1b/c knockdown or increased expression of Homer1a (Tu et al., 1998; Sala et al., 2003; Kammermeier and Worley, 2007; Gasperini et al., 2009). While direct comparison between Homer1b/c^R297W^ and Homer1a was beyond the scope of this study, the convergence of their effects is consistent with a shared dependence on intact tetrameric scaffolding for normal Homer1b/c function. The mechanisms of disruption may differ: Homer1a competes for EVH1-mediated ligand binding with long-form Homer1, whereas Homer1b/c^R297W^ retains its full-length long-form structure but plausibly may compromise scaffolding function by its incorporation into lower-order multimers. That both modes of disruption produce similar phenotypes reinforces the conclusion that the formation of tetrameric scaffolding complexes is the critical determinant of Homer1b/c function.

### Homer1b/c as a regulator of SOCE

Our data provides the first demonstration, to our knowledge, that the integrity of the Homer1b/c coiled-coil domain is required for normal SOCE in neurons. While several studies have implicated Homer1 in SOCE – including observations that Homer1a expression supresses SOCE by disrupting STIM1-Orai1 associations (Jardin et al., 2013; Dionisio et al., 2015; Rao et al., 2016) – prior work had not directly examined the effect of long-Homer knockdown on SOCE. Homer1b/c^R297W^ overexpression significantly attenuated SOCE amplitude in both sensory neuron growth cones and hippocampal neuron somata. In hippocampal neurons, siRNA-mediated Homer1b/c knockdown produced a similar reduction in SOCE, suggesting that Homer1b/c^R297W^ expression and loss of endogenous Homer1b/c converge on the same functional deficit. That Homer1b/c^R297W^ also blunted SOCE in growth cones argues that Homer1b/c scaffolding complexes are necessary for proper scaffolding and regulation of proteins which facilitate SOCE across several neuron types and cellular compartments.

Our super-resolution data may provide a candidate mechanism by which Homer1b/c accomplishes this role. In growth cones, Homer1b/c^R297W^ coupling to STIM1 was significantly reduced, as assessed by both SODA and CBC analysis of dSTORM imaging data. In dendritic spine heads, Homer1b/c^R297W^ expression reduced the density and number of STIM2 clusters present. It is well understood that STIM proteins must translocate to ER-PM junctions and interact with Orai channels to initiate SOCE (Liou et al., 2005; Luik et al., 2006), and Homer1b/c interacts with STIM1 through a PPXXF-compatible proline rich sequence (Dionisio et al., 2015) and promotes the STIM1-Orai1 association that is critical for CRAC channel activation (Rao et al., 2016). We hypothesise that Homer1b/c scaffolding complexes may contribute to the positioning of STIM and ER-PM signalling domains, and that disruption of these scaffolds by Homer1b/c^R297W^ impairs STIM-Orai interactions and hence Ca^2+^ influx through CRACs during SOCE. The reduced density of IP_3_R clusters in Homer1b/c^R297W^ spines, together with the known role of IP_3_R-mediated Ca^2+^ release in triggering SOCE (Thillaiappan et al., 2017), suggests that Homer1b/c^R297W^ may also impair the upstream signalling events that initiate store depletion. Future investigations examining whether Homer1b/c^R297W^ expression directly alters STIM-Orai interactions following ER-Ca^2+^ depletions would help to resolve the precise manner by which these scaffolding disruptions impair SOCE.

### Implications for synaptic plasticity

The scaffolding of group-I mGluRs by Homer1b/c, and the coupling of their activation to downstream ER-Ca^2+^ release via IP3Rs, is among the best characterised functions of long-form Homer1 (Brakeman et al., 1997; Tu et al., 1998; Kammermeier and Worley, 2007). We observed that expression of Homer1b/c^R297W^ disrupted this signalling axis at multiple levels: mGluR5 recruitment to dendritic spines was reduced, mGluR5-Homer1b/c coupling in growth cones was decreased, and the Ca^2+^rise elicited by group-I mGluR activation with 3,5-DHPG was significantly blunted. The reduced density of both mGluR5 and IP3R clusters observed with dSTORM in Homer1b/c^R297W^ spine heads may also provide a structural model under which the attenuation of mGluR-mediated Ca^2+^ signalling we observed can be attributed to a widespread impairment in the post-synaptic scaffolding complexes which tightly link mGluR activation to IP3R-mediated Ca^2+^ release.

Our findings intersect with an existing body of work implicating disrupted Homer-mGluR5 scaffolding as a convergent pathological mechanism across multiple genetic backgrounds associated with ASD and related neurodevelopmental conditions. In mouse models of Fragile X syndrome, enhanced mGluR5 mobility and weakened Homer1b/c-mGluR5 interactions are central to the synaptic and behavioural phenotypes observed, and deletion of Homer1a is sufficient to rescue many features of the disorder in such mice by restoring Homer1b/c-mGluR5 coupling (Ronesi et al., 2012; Guo et al., 2016). Similarly, Shank3 knockout models of ASD present with disrupted Homer-mGluR5 scaffolds alongside reduced spine density and more immature spine morphologies (Peça et al., 2011; Wang et al., 2016). Our data demonstrate that a single substitution within the Homer1b/c coiled-coil domain is sufficient to recapitulate features of these phenotypes, including reduced spine density, impaired mGluR5 clustering and attenuated mGluR-dependent Ca^2+^ signalling. The convergence of these findings across distinct genetic profiles spanning Shank3 loss, FMR1 loss and the Homer1b/c^R297W^ substitution, reinforces the view that the integrity of Homer-mGluR5 scaffolding complexes represent an important node in the molecular pathology underlying altered synaptic connectivity.

### Homer1b/c in axon guidance

Homer1b/c is widely expressed during early CNS development (Gasperini and Foa, 2004; Shiraishi et al., 2004; Foa et al., 2005), and perturbation of its scaffolding function causes axon pathfinding errors at stereotypical choice points in vivo, demonstrating that both the EVH1 and coiled-coil domains are necessary for appropriate guidance (Foa et al., 2001). Subsequent work identified the underpinning mechanism as Ca^2+^-dependent: Homer1 knockdown reverses growth cone turning to BDNF from attraction to repulsion by altering the operational state of the CaMKII/CaN molecular switch, abolishing cue-induced Ca^2+^ release and increasing the frequency of spontaneous Ca^2+^ transients through TRPC channels (Gasperini et al., 2009). More recent studies have demonstrated the necessity for proper functioning of SOCE and its constituent signalling components in BDNF-directed guidance (Mitchell et al., 2012; Pavez et al., 2019), raising the question of how Homer1b/c scaffolding supports these processes at the molecular level during guidance. Our findings refine these models by identifying CC domain integrity as a critical structural determinant of Homer1b/c’s role in Ca^2+^-dependent guidance. dSTORM analysis revealed that Homer1b/c^R297W^ is significantly uncoupled from STIM1 within growth cones, providing a nanoscale correlate for the SOCE deficit, as disrupted CC-mediated scaffolding likely impairs the positioning of STIM1 at ER-PM signalling domains necessary for store-operated Ca^2+^ influx through TRPC and Orai channels. As a de novo variant, Homer1b/c^R297W^ would be present throughout critical periods of circuit formation, suggesting that impaired Homer1b/c scaffolding during development could contribute to the connectivity deficits associated with neurodevelopmental conditions.

### Limitations and future directions

Our hypothesis that Homer1b/c^R297W^ undergoes impaired tetramer formation is grounded in the identification of R297 as a residue mediating intermolecular salt bridges at the C1 dimeric interface of the Homer1b/c coiled-coil (Hayashi et al., 2009). While the functional and super-resolution data presented here are consistent with this model, definitive confirmation would require analytical gel filtration of Homer1b/c^R297W^, both alone and co-expressed with wild-type Homer1b/c, to directly quantify the multimerisation state of the variant in vitro.

It is important to note, however, that the present study employed an overexpression paradigm rather than knock-in of the HOMER1^R297W^ variant at endogenous levels. While the dominant-negative effects we observe suggest that HOMER1^R297W^ can disrupt endogenous Homer1b/c function even when co-expressed, the magnitude of these effects in vivo at physiological expression levels remains to be determined. Generation of a CRISPR-knock in model expressing Homer1b/c^R297W^ would enable assessment of the variant’s effects under physiological expression conditions, in the context of human cellular contexts in iPSC-derived neurons, and on circuit development through animal models, which would be necessary to address this question.

The Ca^2+^ signalling deficits and cellular phenotypes we report are likely to be mechanistically linked, given the well-established dependence of processes including both spine maturation and axon guidance on tightly regulated Ca^2+^ dynamics. However, we have not directly demonstrated here a causal relationship in the context of Homer1b/c^R297W^-induced phenotypes. Future experiments in which SOCE or mGluR-dependent Ca^2+^ signalling is pharmacologically rescued in Homer1b/c^R297W^-expressing neurons would help to establish whether restoring normal Ca^2+^ dynamics is sufficient to ameliorate the various molecular and cellular phenotypes we report here and may expand our understanding of the molecular processes by which Homer1b/c performs such important regulatory influence over various forms of synaptic functioning.

## Conclusion

Here we identify *HOMER1^R297W^* as a de novo variant that disrupts Homer1b/c scaffolding through a single substitution within the coiled-coil domain. Expression of Homer1b/c^R297W^ attenuated Ca²⁺ signalling in sensory neuron growth cones and hippocampal neurons, impaired both axon guidance and spine density and reduced the clustering of Ca²⁺ handling proteins within axonal and dendritic compartments. These findings position coiled-coil domain integrity as a critical determinant of Homer1b/c functions and highlight the broader importance of intact scaffolding complexes in coordinating the molecular processes which underpin neuronal development, plasticity and connectivity.

## Author contributions

D.B., L.F., and R.G designed research; D.B. performed research; D.B wrote the manuscript.

